# Single cell transcriptomics of human skin equivalent organoids

**DOI:** 10.1101/2022.07.27.501753

**Authors:** Adam R. Stabell, Shuxiong Wang, Grace E. Lee, Ji Ling, Sandrine D. Nguyen, George L. Sen, Qing Nie, Scott X. Atwood

**Author notes:** Corresponding author: Scott X. Atwood.

## Abstract

Several methods for generating human skin equivalent (HSE) organoid cultures are regularly used to study skin biology and test pharmaceuticals, however few studies have thoroughly characterized these systems. To fill this gap, we used single cell-RNA sequencing to compare the cellular states of *in vitro* HSEs generated from distinct culture methods, HSEs xenografted onto mice, and *in vivo* epidermis. By combining differential gene expression, pseudotime analyses, splicing kinetics, and spatial localization, we reconstructed HSE keratinocyte differentiation trajectories that recapitulated known *in vivo* epidermal differentiation pathways and show that HSEs contain many of the major *in vivo* cellular states. However, HSEs also develop several unique keratinocyte states, an expanded basal stem cell program, and disrupted terminal differentiation. In addition, cell-cell communication modeling showed the presence of EMT-associated signaling pathways not normally active in homeostatic skin and we show that EGF supplementation influences the EMT signature. Lastly, xenografted HSEs at early timepoints post-transplantation significantly rescued many of the observed *in vitro* deficits, while undergoing a hypoxic response that drove an alternative differentiation lineage. This study highlights the strengths and limitations of organoid cultures and identifies areas for potential innovation.

## INTRODUCTION

Skin is an essential organ with many roles including forming a water-tight barrier, aiding in thermoregulation, and acting as a sensory organ (Kolarsick et al., 2011). To fulfill these roles, the keratinocytes that constitute the epidermis must replenish themselves while withstanding a constant barrage of chemical, physical, pathological, and radiological insults from their environment (Ghazizadeh and Taichman, 2005; Khavkin and Ellis, 2011). The field of skin research has largely been driven by *in vivo* mouse models that show healthy skin is critical to an organism’s wellbeing and the disruption of many of its functions can lead to a drastic decline in quality of life (Corrò et al., 2020; Zomer and Trentin, 2018). While mice are suitable to define the basic architecture and homeostatic signaling of skin, the anatomy, microstructure, and heterogeneity of mouse skin is inherently different from human (Kolarsick et al., 2011). For instance, mice have a distinct density of hair follicles and eccrine glands, a layer of striated muscle found beneath the hypodermis, a lack of melanocytes in the interfollicular epidermis, and the absence of rete ridges. These differences impact epidermal homeostasis, wound repair, and the severity of certain skin disorders, pointing to a need for a more human equivalent model system to study human-specific aspects of skin biology (Zomer and Trentin, 2018).

Three-Dimensional (3D) organoid cultures have long been a tool to investigate complex tissue interactions. Typically composed of primary cells isolated from patient samples, the idea of building an organ from its basic components is an attractive premise that has profound scientific implications (Atwood and Plikus, 2021). From gaining molecular insight by simplifying development and homeostasis to their essential parameters to the translational promise of a gold standard system to test drugs or a farm system to grow replacement tissues, 3D organoid cultures are gaining popularity as an elegant and relevant model system to study human biology. Current technologies include generating complex skin in spherical cell aggregates from human pluripotent stem cells (Lee et al., 2020; Zhang et al., 2021), using conventional scaffolds - such as hydrogels (Fauzi et al., 2020; Morrison et al., 2020; Stark et al., 1999) or bioprinting (Patra and Young, 2016; Pourchet et al., 2017; Zhang et al., 2021) - to assemble dermal and keratinocyte layers with other relevant cells, and organ-on-a-chip that allows active perfusion and spatiotemporal control at the microscale level (Zhang et al., 2018).

However, 3D cultures are not without their limitations. For instance, despite human skin equivalent (HSE) organoid cultures showing a high degree of morphological similarity to their *in vivo* counterparts, their composition and culturing conditions vary greatly from lab to lab which can affect interpretation of similar experiments (Atwood and Plikus, 2021; El Ghalbzouri et al., 2009; Lee et al., 2020; Li and Sen, 2015; Pourchet et al., 2017; Zhang et al., 2021). Many components of the *in vivo* system are lacking, such as vasculature and immune cells, which limit the size of cultures and their response to experimental stimuli (Atwood and Plikus, 2021). And many studies defining HSEs have shown marked molecular differences in basal and terminal gene expression that suggest epidermal differentiation is not quite analogous to their *in vivo* counterparts (Andreadis et al., 2001; Jevtić et al., 2020). Given the variability that exists between culture systems and their limited characterization, it can be difficult to determine which conditions are best suited for a particular experiment. Knowledge of the capacity and limitations of these systems is paramount to accurately interpret their results.

Recently, several labs have published single cell -omic studies examining the strengths and weaknesses of a variety of organotypic culture systems. These include organoids mimicking the central nervous system (Brancati et al., 2020), gastric system (Chen et al., 2019), intestinal system (Serra et al., 2019), and gastrulation (van den Brink et al., 2020). Human skin spheroids have recently been developed from human pluripotent stem cells that differentiate into spherical cell aggregates where cyst-like skin emerges composed of stratified epidermis, fat-rich dermis, pigmented hair follicles with sebaceous glands, and rudimentary neural circuitry (Lee et al., 2020). Although these skin spheroids resemble fetal facial skin, their long incubation period and small size are not ideal for genetic manipulation of individual cell types or for grafting in the clinic. How HSEs built using conventional scaffolds like devitalized dermis compare to their *in vivo* counterparts is unclear, despite being ideally suited to address the deficiencies of spherical skin organoids.

Our lab, alongside others, have recently shown that human epidermis is more heterogeneous than previously thought (Cheng et al., 2018; Reynolds et al., 2021; Solé-Boldo et al., 2020; Wang et al., 2020b). Using single cell-RNA-sequencing (scRNA-seq) and subsequent *in vivo* validation, we spatially resolved four distinct basal stem cell populations within human interfollicular epidermis and delineated multiple spinous and granular cell populations that contributed to a hierarchical differentiation lineage supporting multi-stem cell epidermal homeostasis models (Wang et al., 2020b). Collectively, these studies have highlighted the complexity of the epidermis and their cell-cell interactions. The extent to which HSEs are capable of recapitulating the cell type heterogeneity, cell-cell signaling, and differential gene expression of *in vivo* human skin remains unclear. To address this issue, we probed the transcriptomes of three HSE variants and examined the differences in comparison to *in vivo* human skin at the single cell level. We found that all HSEs remarkably contained the relevant cellular states of their *in vivo* counterparts, but each HSE also possessed unique cell states not found during homeostasis. An expanded basal program, terminal differentiation defects, and ectopic EMT signatures predominate fibroblast- and Matrigel-derived HSEs, whereas xenografting HSEs onto immunodeficient mice largely rescued the various defects at the cost of inducing hypoxic conditions.

## RESULTS

### Histological characterization of HSEs

To compare commonly used *in vitro* HSEs to *in vivo* human epidermis, we chose to use devitalized human dermis as the scaffold for growing the HSEs because we reasoned that the extracellular matrix composition more accurately mimics the endogenous surface for keratinocyte stratification than a collagen-based hydrogel. We utilized the two most common HSE variants where primary human keratinocytes are seeded on top of devitalized dermis at the air-liquid interface and the dermis is either treated with Matrigel (GelHSE) or seeded with primary human dermal fibroblasts (FibHSE) to supply necessary signals for keratinocyte stratification (**Figure 1A**). Keratinocyte stratification occurs under both conditions by day 7, where the HSEs show a tightly packed columnar basal cell layer, multiple irregular polyhedral squamous cell layers, several flattened granular cell layers, and a thin stratum corneum (**Figure 1B**). Histologically, the HSEs largely remain the same up through day 28 except for a thickening of the stratum corneum and a general spreading out of keratinocytes at all epidermal layers. Proliferation was also reduced in the HSEs compared to neonatal or adult epidermis (**Figure 1C-D**). FibHSEs possess a significantly thicker living epidermal layer than the GelHSEs (**Figure 1E**). We chose to continue our analysis with day 28 HSEs due to the morphological similarity to *in vivo* tissue and to avoid active re-stratification programs that may be operating at earlier timepoints.

**Figure 1:**
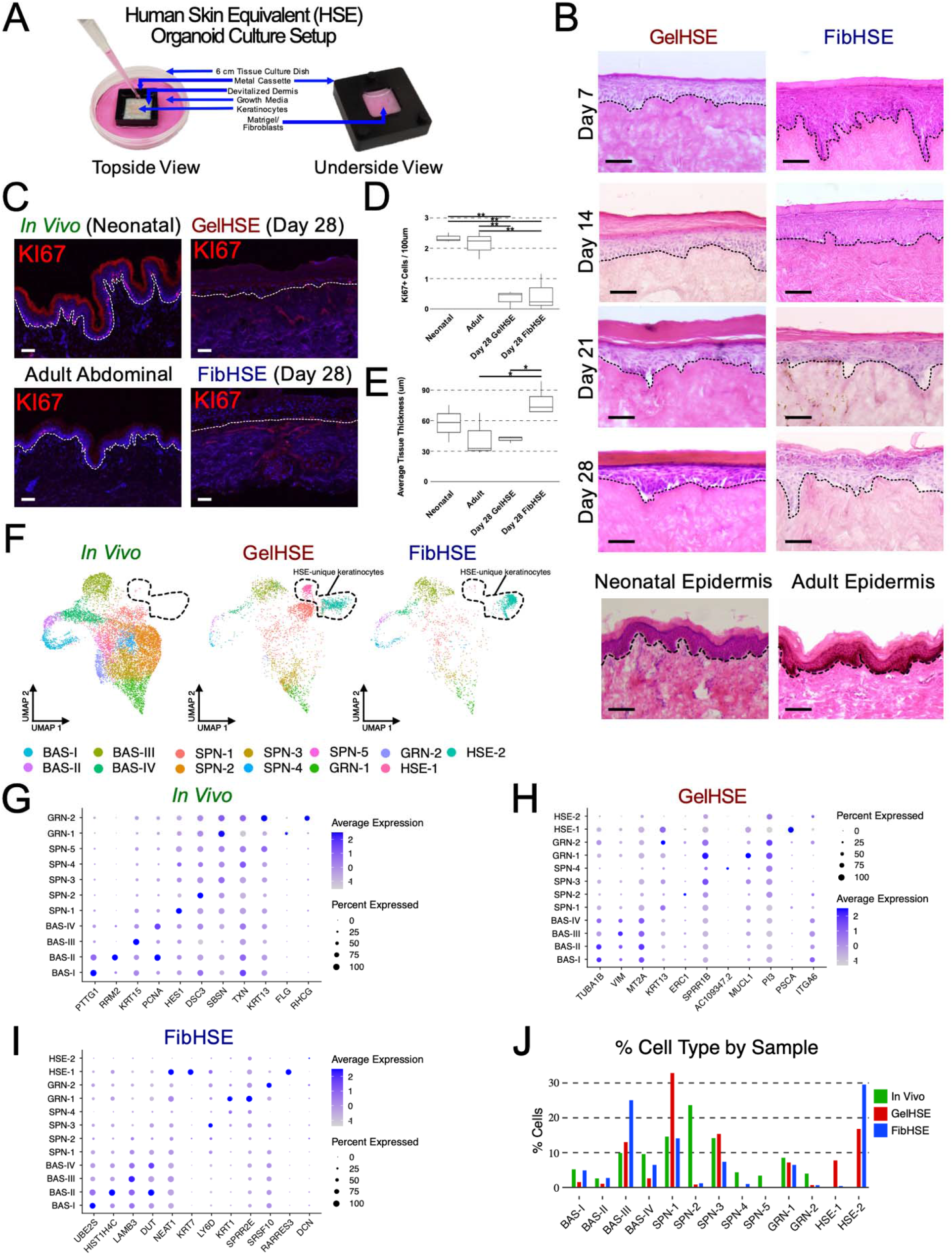
Defining human skin equivalent cell populations using scRNA-seq. **A)** Diagram of the human skin equivalent (HSE) organoid culture setup. **B)** Hematoxylin and eosin (H&E) staining of Matrigel-grown HSEs (GelHSE) and fibroblast-seeded HSEs (FibHSE) after 7, 14, 21, and 28 days of growth on devitalized human dermis. Neonatal epidermis from foreskin and adult epidermis from the leg are shown for comparison. Scale bar 100 μm. Dashed lines denote the epidermal-dermal junction. **C)** Immunostaining of KI67 (red) and DAPI (blue) in human neonatal skin (top left), adult abdominal skin (bottom left), day 28 GelHSE (top right), and day 28 FibHSE (bottom right). Scale bar 100 μm. Dashed lines denote the epidermal-dermal junction. Quantification of **(D)** KI67+ cells and **(E)** average thickness of living epidermal cell layers in human neonatal skin, adult abdominal skin, day 28 GelHSE, and day 28 FibHSE. n = 3 each sample. Significance was determined by Tukey’s HSD test. *p < 0.05. n.s., not significant. **F)** Seurat clustering of 16,791 single cells isolated from four HSE libraries (two GelHSE and two FibHSE) and two *in vivo* neonatal epidermis libraries using UMAP embedding. Libraries are split by sample type. Dashed lines encompass HSE-unique keratinocytes. Dot plots of the top differentially expressed marker genes for **(G)** *in vivo* clusters, **(H)** GelHSE clusters, and **(I)** FibHSE clusters. **J)** Percentage of total cells within each cluster split by sample type.

### Epidermal homoeostasis is disrupted in HSEs

To define the cellular states of keratinocytes derived from HSEs, we isolated viable, single cells from day 28 HSEs and subjected them to droplet-enabled scRNA-seq to resolve their individual transcriptomes (**Supplemental Figure 1A**). We processed a total of 4,680 cells from two FibHSEs (including fibroblasts) and 4172 cells from two GelHSEs before performing quality control analysis on individual libraries using the R package Seurat (**Supplemental Figure 1B**). The cells from each replicate FibHSE were clustered in an unsupervised manner, and tentatively annotated as keratinocytes or fibroblasts using the marker genes *KRT14* and *KRT10* to identify keratinocytes and *TWIST2* and *COL6A1* to identify fibroblasts (**Supplemental Figure 2**). Keratinocytes were then subset from our HSE datasets and integrated with interfollicular keratinocytes from two *in vivo* human neonatal epidermal datasets that were previously generated by our lab (Wang et al., 2020b)(**Figure 1F**). Cell types were annotated based on known marker genes from the *in vivo* dataset, which differed from the marker genes of the HSE datasets (**Figure 1G-I**). Remarkably, many of the major *in vivo* cellular states were found in the *in vitro* HSEs, including the full complement of *in vivo* basal cell states. Based on our previous characterization of basal stem cell communities (Wang et al., 2020b), BAS-I – BAS-IV represented approximately 27.3% of the *in vivo* cells, 39.2% of FibHSE, and 18.3% of GelHSE, and were enriched for known basal keratinocyte marker genes including *PTTG1, RRM2, KRT15,* and *PCNA,* respectively (**Figure 1G-I**). The ratios of BAS-I and BAS-II cycling cells remained largely similar, despite a reduction in cycling cells in the GelHSE, whereas BAS-III cells are enriched, and BAS-IV cells depleted in the HSEs compared to their *in vivo* counterpart (**Figure 1J**). Intriguingly, two of the HSE cell states clustered separately from the *in vivo* cells and were annotated HSE-1 and HSE-2 (**Figure 1F**). HSE-specific keratinocytes constituted 29.6% of FibHSE and 24.6% of GelHSE (**Figure 1J**). HSE-1 is unique to GelHSE, whereas HSE-2 appears in both HSE samples with a greater prevalence in FibHSE (**Figure 1J**).

Despite the relatively normal histological appearance of the HSEs, there is an expansion of KRT14+ cell layers and disrupted epidermal differentiation in both the GelHSE and FibHSE cultures (**Figure 2A, Supplemental Figure 3**). The expanded KRT14+ cell layers do not proliferate outside of the basal layer in contact with the basement membrane (**Figure 1C**) and differentiation markers such as DSG1, FLG, and LOR are still restricted from the basal-most layer (**Figure 2A-B**). Basal cell marker KRT15 does remain restricted to the basal-most layer of the HSEs, whereas KRT19 shows selective expansion in the GelHSE (**Figure 2C**), suggesting that suprabasal KRT14+ cells are not fully functioning basal cells and are likely to be differentiating without fully turning off the basal cell state. The mesenchymal marker VIM, which is normally restricted to fibroblasts, melanocytes, and Langerhans cells of *in vivo* skin, shows high RNA expression in GelHSE basal keratinocytes and VIM+ protein expression in both GelHSE and FibHSE basal keratinocytes (**Figure 2D**), suggesting a partial epithelial-to-mesenchymal transition (EMT) state. This partial EMT state is not entirely unexpected given the signals the keratinocytes are receiving from the Matrigel and culture media, with the GelHSE showing the greatest expression of *VIM.* Cell-cell contacts and terminal differentiation are also disrupted in HSEs with DSG1 protein no longer restricted to cell-cell contact sites, FLG protein expression turning on early in SPN cell layers, and FLG and LOR no longer restricted to the GRN layers (**Figure 2A-B**) Despite testing several marker genes, we were unable to spatially resolve HSE-2. However, HSE-1 is readily identified by one of its marker genes, *PSCA* (**Figure 2D**). *PSCA* encodes for a GPI-anchored membrane glycoprotein typically found in basal cells of the prostate, the lining of the urinary tract, the mucosal epithelium of the gastrointestinal tract, and in the outermost layer of fetal skin from E15 to E17 (Ross et al., 2001). Staining for PSCA demonstrated that these keratinocytes are exclusively localized to the outermost epidermal layers (**Figure 2D**) and may indicate a remnant embryonic program that is reactivated as a result of growth factors in the culturing media.

**Figure 2:**
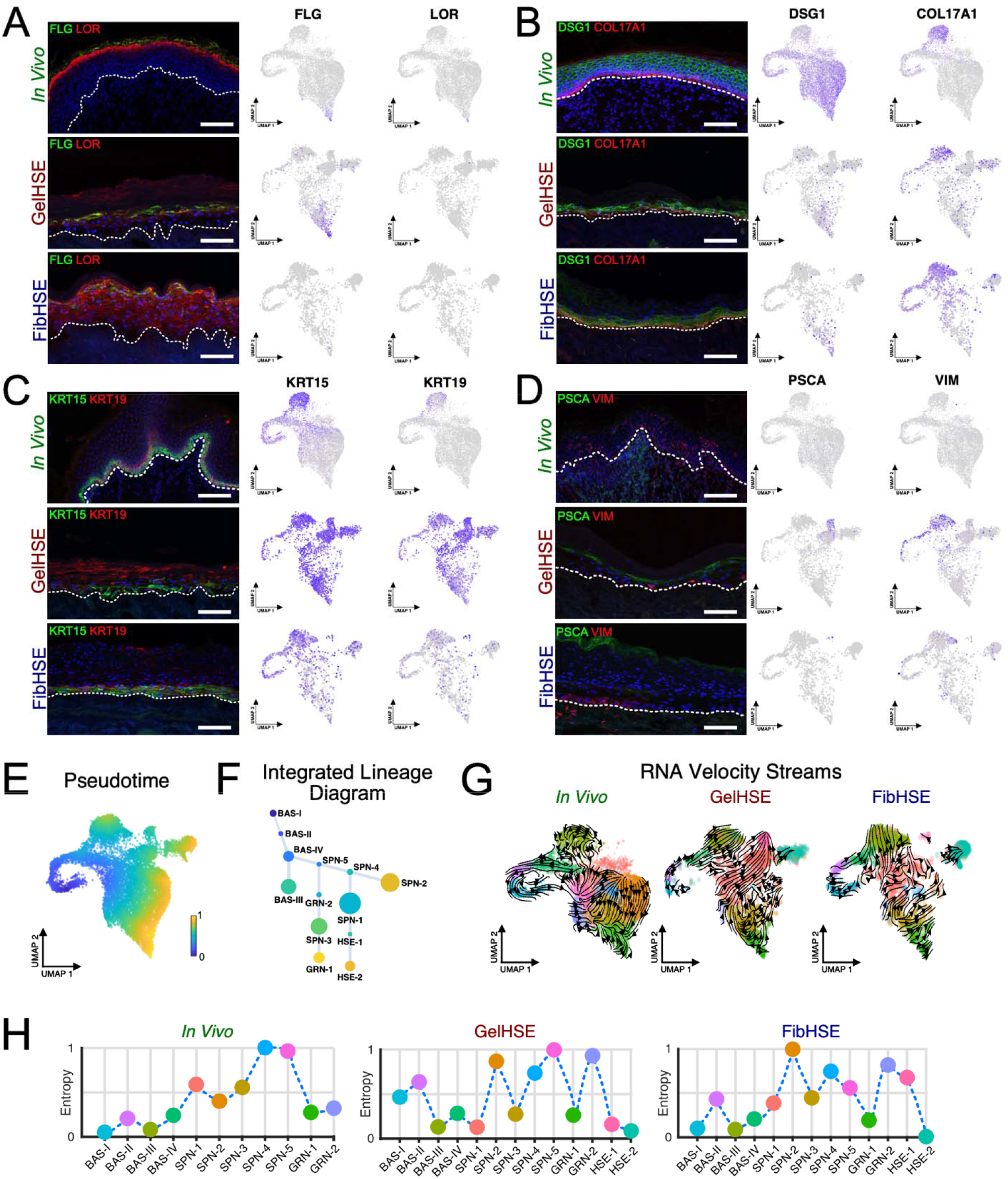
Human skin equivalents display altered expression patterns and lineage paths. Immunostaining of **(A)** terminal differentiation markers FLG and LOR, **(B)** structural proteins DSG1 and COL17A1, **(C)** basal stem cell markers KRT15 and KRT19, and **(D)** human skin equivalent (HSE) unique markers PSCA and VIM. Human neonatal skin (top), day 28 GelHSE (middle), and day 28 FibHSE (bottom). Feature plots showing the RNA expression of indicated markers for each sample type are on the right. Scale bar 100 μm. Dashed lines denote the epidermal-dermal junction. **E)** Pseudotime inference of epidermal keratinocytes from the integrated datasets. **F)** Cell lineage diagram of keratinocytes from the integrated datasets. Edge weights denote probability of transition to each cluster. Dot size denotes number of cells. **G)** Splicing kinetics depicted as RNA velocity streams calculated using the python package scVelo. **H)** Quantification of Cellular Entropy (*ξ*) using the R package SoptSC.

Considering the apparent uncoupling of markers from their respective cell states, we averaged the RNA expression of every cell in each cluster and calculated a Pearson correlation between the HSE and *in vivo* clusters (**Supplemental Figure 4A**). Both *in vivo* datasets were compared to each other to establish the highest expected Pearson correlation between cell states. With respect to the HSEs, the most highly correlated clusters were the basal cell populations. Interestingly, the majority of HSE clusters showed the highest correlation with the *in vivo* BAS-III cluster, suggesting that the BAS-III transcriptional program is not shut off during HSE differentiation. Additionally, the Pearson correlation decreases as keratinocytes differentiate, reinforcing that terminal differentiation is disrupted in HSEs. The correlation between the *in vivo* tissue and FibHSE is higher overall than GelHSE, indicating that global RNA expression in FibHSE more accurately mimics *in vivo* human epidermis.

### Human skin equivalents have altered lineage paths

Next, we examined how the HSE-specific clusters altered the inferred lineage trajectory of epidermal differentiation. We generated pseudotime and cell lineage inferences of the integrated keratinocytes using SoptSC (Wang et al., 2019) and partially reconstructed the expected BAS-SPN-GRN keratinocyte differentiation trajectory (**Figure 2E-F**). Basal keratinocytes expressing *KRT15* were placed at the beginning of the trajectory and cells expressing the terminal differentiation gene *FLG* were placed towards one of the trajectory termini (**Figure 2F**). Intriguingly, the HSE-specific clusters were placed at a distinct trajectory terminus away from the GRN cell states, with HSE-1 preceding the more differentiated HSE-2 and generating a BAS-SPN-HSE differentiation trajectory (**Figure 2F**).

To better define the BAS-SPN-HSE differentiation trajectory, we analyzed the splicing kinetics of every cell using scVelo’s dynamical modeling, to infer the future state of each cluster (Bergen et al., 2020). We subset cells from each tissue from the integrated dataset and modelled them separately while keeping their spatial relationship within the integrated dataset intact (**Figure 2G**). The *in vivo* epidermal dataset showed the expected BAS-III and BAS-IV velocity vectors pointing towards the SPN clusters and SPN velocity vectors pointing towards the GRN clusters, reconstructing the BAS-SPN-GRN differentiation trajectory (**Figure 2G**). While the FibHSE trajectory largely followed this trend, many BAS and SPN velocity vectors for both HSEs point towards the HSE-1 and HSE-2 clusters, with an undefined flow of vectors between the SPN and GRN clusters, suggesting that terminal differentiation may be disrupted and that the HSE clusters may represent an alternative differentiation trajectory terminus.

We next used SoptSC’s cellular entropy estimator to infer the entropy of each cluster to determine the relative stability of each cellular state (Wang et al., 2020b). High entropy suggests a high probability that a cell will transition into another state and low entropy indicates a low probability that a cell will transition into another state. The *in vivo* epidermal dataset shows low entropy for the BAS and GRN clusters, indicating that these are stable states, whereas the SPN clusters have high probabilities of transitioning to a new state (**Figure 2H**). These *in vivo* entropy values reinforce the idea that once differentiation is initiated in the SPN state, there is momentum to reach terminal differentiation in the GRN state as an endpoint with high energy costs to stop at any intermediate stage. For the GelHSE and FibHSE datasets, BAS-III, BAS-IV, and GRN-1 remain stable states, suggesting that these states are robust to perturbations and remain a core lineage trajectory in the HSEs (**Figure 2H**). However, the HSE-2 state has the lowest entropy values out of all the other epidermal states, suggesting this stable state is a likely alternative endpoint and reinforcing the BAS-SPN-HSE differentiation trajectory as an alternative trajectory to BAS-SPN-GRN in HSE cultures.

### HSEs exhibit abnormal signaling associated with EMT

We sought to infer how intercellular communication is altered in the HSEs using CellChat, a bioinformatic tool that predicts intercellular communication networks using ligands, receptors, and their cofactors to represent known heteromeric molecular complexes instead of the standard one ligand/one receptor gene pair (Jin et al., 2021). CellChat detected 17 significant signaling pathways in the *in vivo* dataset and the HSEs recapitulated 14 of the 17 pathways (**Supplemental Figure 4B, Supplemental Tables 1-3**). However, the HSEs also showed an extended network of significant signaling pathways, with 34 in GelHSE and 33 in FibHSE. A subset of these pathways, such as LAMININ, CD99, CDH1, EPHB, and MPZ, show similar signaling profiles across the *in vivo* and HSE tissues, whereas the other pathways show marked differences (**Supplemental Figure 4B**). Many of the outgoing and incoming signals in the *in vivo* dataset predominantly come from or go to the BAS-III and GRN-1 clusters, suggesting that these stable cell states have great influence over tissue function (**Supplemental Figure 4B**). While BAS-III and GRN-1 are still signaling hubs in the GelHSE and FibHSE datasets, HSE-1-specific signaling exerts wide influence over GelHSE whereas all four BAS clusters actively signal in FibHSE with little contribution to or from HSE-specific clusters.

Given the abnormal VIM expression in the HSE basal keratinocytes that is normally found in mesenchymal cells, we decided to explore EMT signaling in the HSEs. We focused on EGF signaling, a well-documented inducer of EMT (Kim et al., 2016). EGF signaling in *in vivo* epidermis mainly comes from the differentiated GRN or more differentiated SPN cell populations and signals to the BAS stem cell and early SPN populations (**Figure 3A**). However, sender EGF signaling is expanded to the BAS and early SPN populations in the HSEs, coinciding with the appearance of VIM+ basal cells (**Figure 3A**). Both HSE-unique clusters are involved in EGF signaling, with HSE-2 sending signals in both HSE cultures and HSE-1 sending and receiving signals in GelHSE. This prominent role in EGF signaling is unusual for HSE-2, which plays a very small role in global signaling events (**Supplemental Figure 4B**). Interestingly, the ligands and receptors facilitating EGF signaling are substantially altered in both HSEs compared to the *in vivo* state (**Figure 3B-C**). AREG-EGFR signaling is overrepresented in both HSEs and the *AREG* ligand is expressed in most HSE-cultured keratinocytes (**Figure 3B-C**). *EREG* and *TGFA* ligands also specifically contributes to EGF signaling in the HSEs, whereas HBEFG-EGFR signaling is reduced compared to the *in vivo* state (**Figure 3B-C**). These ligands have all been implicated in EMT induction by activation of the EGFR/ERK/NF-κB signaling pathway (Aharonov et al., 2020; Eapen et al., 2019; Liu et al., 2016; Shostak et al., 2014; Wang et al., 2020a; Yu et al., 2018).

**Figure 3.**
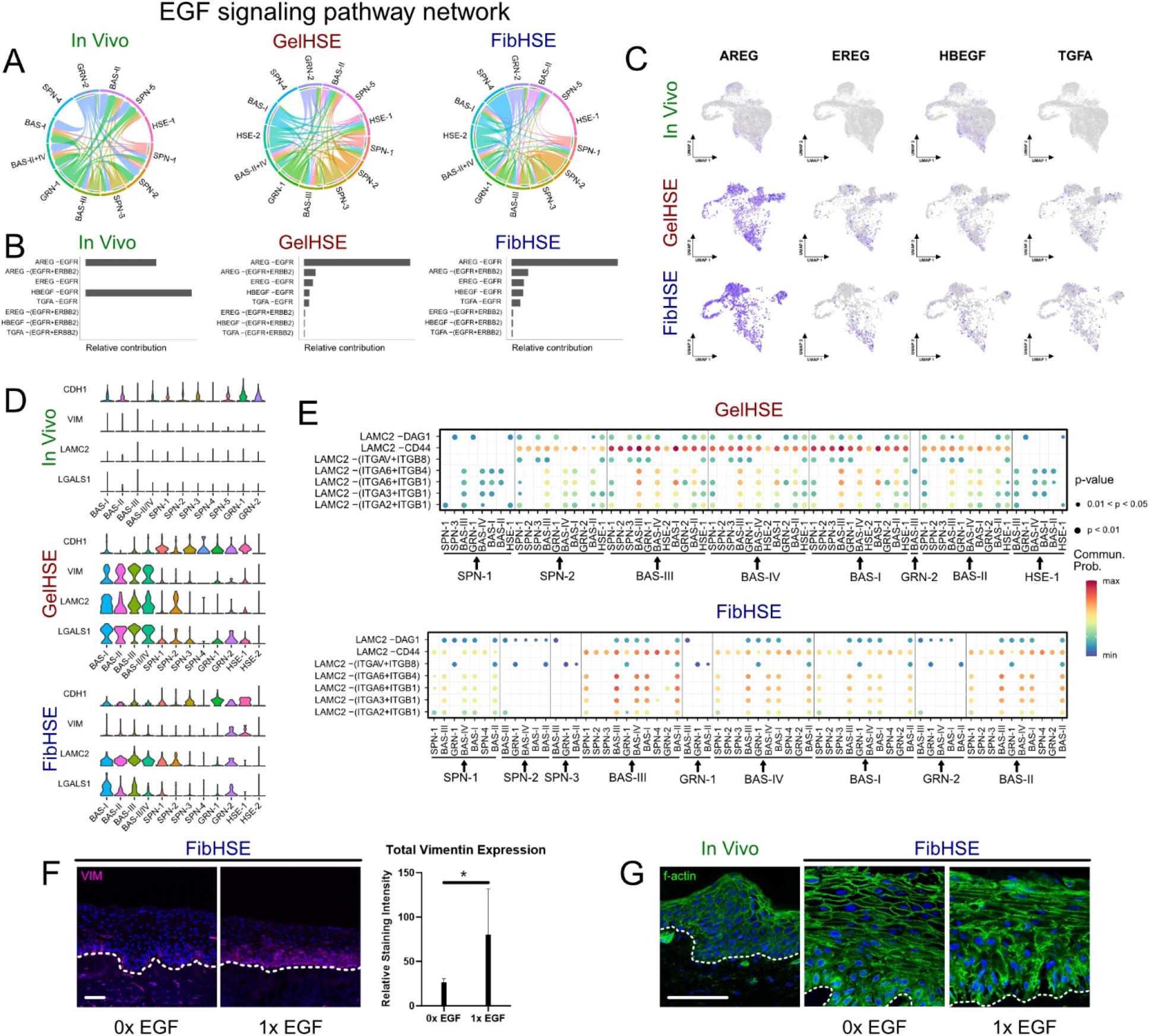
HSEs possess an EMT-like gene expression signature driven by EGF sigaling. **A)** Cell-cell communication networks predicted for the EGF signaling pathway inferred using the R package CellChat. Edge weights represent the probability of signaling between cell clusters. **B)** Relative contributions of each ligand, receptor, and cofactor group to the cell-cell communication predicted in panel A. **C)** Feature plots showing the expression patterns of each of the ligands contributing to the EGF signaling network. **D)** Violin plots of relative gene expression for positive markers *(VIM, LAMC2,* and *LGALS1)* and negative markers *(CDH1)* of EMT. **E)** Visualization of signaling probability scores of ligand-receptor/co-receptor pairs involving *LAMC2* for GelHSE and FibHSE datasets. *In vivo* datasets had no imputed signaling interactions involving LAMC2. Dot size represents p-value. **F)** Immunostaining of VIM in FibHSEs supplemented with indicated concentrations of EGF. Quantification of VIM staining intensity is shown on the right. n = 3 each condition. One-tail student’s t-test was used to determine significance. * denotes p-value < 0.1. Scale bar = 100 μm. **G)** Filamentous actin (factin) staining of FibHSEs supplemented with indicated concentrations of EGF. Scale bar = 100 μm.

Several other genes associated with EMT, such as *LAMC2* and *LGALS,* are also expressed in HSEs (**Figure 3D**). LAMC2 is a regulator of the EMT phenotype and silencing LAMC2 reverses EMT by inactivating EGF signaling (Okada et al., 2021; Pei et al., 2019), whereas *LGALS1* promotes EMT and may be a biomarker of this process (Bacigalupo et al., 2015; Li et al., 2017). Both HSE cultures have high levels of *LAMC2* and *LGALS1* expression in all basal populations, and lower expression levels in more differentiated keratinocytes (**Figure 3D**), supporting the notion that many of the HSE basal cells may be undergoing EMT. *VIM, LAMC2,* and *LGALS1* expression are all higher in the Matrigel-supported GelHSE compared to the FibHSE cultures. Epithelial cell marker *CDH1* is negatively correlated with *VIM* and shows higher expression in *VIM-* HSE keratinocytes compared to the *in vivo* state (**Figure 3D**), suggesting that VIM+ keratinocytes may lose contact with the underlying basement membrane and potentially explaining the small gaps we observe between basal keratinocytes and the basement membrane in older HSE cultures (**Figure 1B**). Furthermore, *LAMC2* shows high probability interactions with several integrins expressed in basal keratinocytes, including *ITGA6, ITGB4, ITGB1,* and the cell-surface glycoprotein *CD44* (**Figure 3E**). *CD44* undergoes complex alternative splicing and at least one of these isoforms is implicated in EMT (Miyazaki et al., 2018; Xu et al., 2015).

To define the relationship between EGF signaling and VIM+ basal cells, FibHSEs were grown in normal growth media for one week to induce epidermal stratification and then the media was supplemented with either 0x, 1x, 2x, or 4x EGF for an additional week (**Figure 3F-G and Supplemental Figure 5**). Removal of EGF resulted in a significant decrease in VIM expression in FibHSE keratinocytes (**Figure 3F**), whereas further EGF supplementation increased VIM expression (**Supplemental Figure 5**). Intriguingly, the nuclei of FibHSE keratinocytes cultured without EGF appeared more ordered and morphologically similar to *in vivo* basal keratinocytes. To characterize the cytoskeletal rearrangements that may be underlying the morphology changes, we stained for filamentous actin and observed qualitative changes in the 0x EGF cultured FibHSEs that more resembled the *in vivo* state (**Figure 3G**). These data suggest EGF supplementation may be a major driver of EMT in the HSE organoid cultures.

### Xenografting partially rescues HSE abnormalities

Despite using devitalized human dermis as a substrate, HSE organoid cultures have a simplified cellular composition that lack system-level aspects of normal skin, such as fully functioning vasculature, immune system, and innervation. One way to circumvent some of these issues is to xenograft HSE cultures onto mice to more accurately mimic endogenous conditions. To explore how the cellular states and transcriptional profile of HSEs were altered when xenografted onto mice, we grew three GelHSE cultures for one week and subsequently grafted them onto a wound bed created within the dorsal back skin of NOD-SCID gamma (NSG) mice where they remained for 24 additional days before dissecting the tissue for scRNA-seq (**Figure 4A-B**). NSG mice were chosen due to their ability to engraft skin at very high levels and perivascular infiltration of immune cells (Brehm and Shultz, 2012). Cell suspensions from the three GelHSE cultures were pooled prior to sequencing. The xenograft dataset was aligned and annotated twice, once using the human reference genome GRCh38 and again using the mouse genome mm10. Mitochondrial gene expression and RNA features were used to identify mouse and human cells (**Supplemental Figure 6A-B**). Human cells have more nuclear and mitochondrial RNA reads aligning to a human reference genome, and the same is true for mouse reads and a mouse reference genome (**Supplemental Figure 6C-D**). After regressing out mouse cells, the dataset was compared to the *in vivo* epidermal datasets in the same manner as our HSE analyses. Surprisingly, we observe four xenograft-unique clusters in the xenograft, alongside the expected BAS, SPN, and GRN keratinocyte clusters (**Figure 4C**).

**Figure 4.**
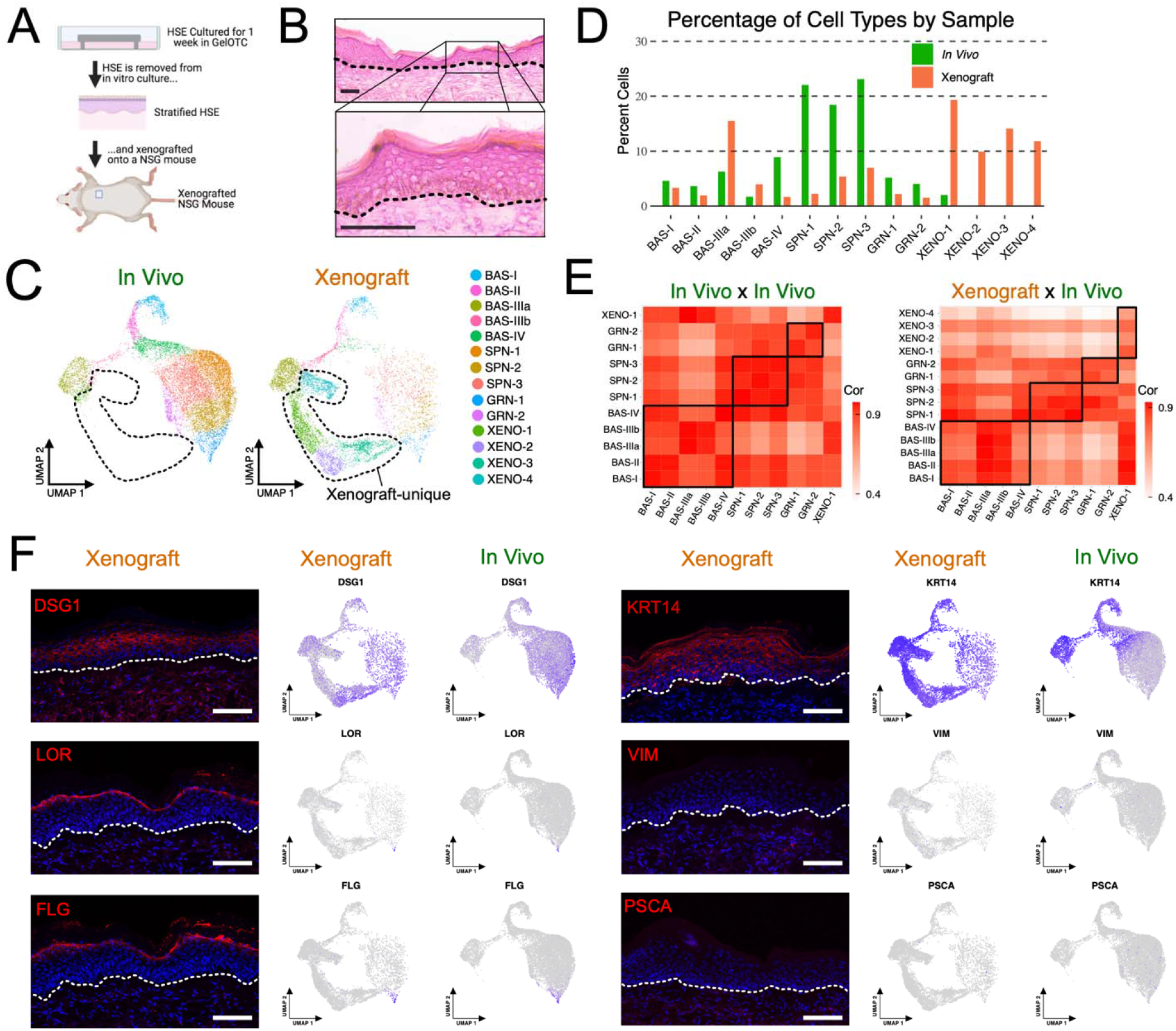
Xenografting rescues terminal differentiation, cell-cell adhesion, and organoid-specific programs. **A)** Schematic of strategy to xenograft human skin equivalent (HSE) tissue. **B)** H&E staining of xenograft tissue. Scale bar 100 μm. Dashed lines denote the epidermal-dermal junction. **C)** Seurat clustering of single cells isolated from pooled xenograft libraries (n = 3 samples pooled prior to sequencing) and two neonatal epidermal libraries and displayed using UMAP embedding. Libraries are split by sample type. Dashed lines encompass xenograft-unique clusters. **D)** Percentage of total cells within each cluster split by sample type. **E)** Pearson correlation of average RNA expression of each cluster compared to all other clusters between the *in vivo* datasets (left) and between the xenograft dataset and both *in vivo* datasets (right). **F)** Immunostaining of indicated markers in HSE xenografted tissue. Feature plots showing the RNA expression of indicated markers are to the right. Scale bar 100 μm. Dashed lines denote the epidermal-dermal junction.

The four xenograft-unique clusters were designated XENO-1 through XENO-4 and collectively comprise ~55% of the total xenograft cells (**Figure 4D**). To better define the difference between the HSE and XENO cellular states, we subset and integrated the HSE-unique cells (HSE-1 and HSE-2) with the xenograft-unique cells (XENO-1 through XENO-4). The xenograft-unique keratinocytes cluster separately from the HSE-unique cells (**Supplemental Figure 6E)**, suggesting that the HSE-specific keratinocytes are unique to organoid culturing and that the xenograft-unique keratinocytes are new cellular states induced after engraftment.

All of the *in vivo* cellular states are present in the xenograft HSEs (**Figure 4C-D**). However, the proportion of BAS-III and BAS-IV keratinocytes are not similar to each other, with BAS-III proportions being much higher and BAS-IV being lower in the xenograft than the *in vivo* setting (**Figure 4D**), a relationship found in the GelHSE and FibHSE cultures (**Figure 1H**) and suggesting that the abnormal basal cell proportions are not rescued by engraftment. The correlation between *in vivo* cell states improves in the xenograft cultures compared to the HSE cultures and the BAS-III state is no longer expanded into the SPN and GRN states (**Figure 4E vs Supplemental Figure 6**). Histologically, the xenografts appear relatively normal, with some basal keratinocytes adopting a cuboidal morphology (**Figure 4B**). Terminal differentiation appears to be rescued as RNA expression and immunofluorescence staining of FLG and LOR are now restricted to the granular layer and cell-cell contacts appear more normal with DSG1 now localizing to cell-cell contact sites (**Figure 4F**), suggesting that barrier formation, which is disrupted in HSE cultures, may be rescued upon engraftment. The basal cell states still appear to be partly disrupted, where total RNA expression for all four BAS clusters in the xenograft have the highest correlation with *in vivo* BAS-III rather than their respective cluster (**Figure 4E**), and KRT14 protein and RNA are still expanded into suprabasal layers (**Figure 4F**). Several basal cell markers are now appropriately expressed in their corresponding cell states compared to the HSE cultures *(PTTG1* with BAS-I, *RRM2* with BAS-II, and *ASS1* and *KRT19* with BAS-III), with *COL17A1* still showing abnormal expression (**Supplemental Figure 7**). The two abnormal features of the HSE cultures, the partial VIM+ EMT-like state and remnant PSCA+ embryonic program, are no longer detected in the xenograft tissue (**Figure 4F**), suggesting that the two abnormal programs seen in the HSEs are rescued. The xenograft-unique clusters notwithstanding, the xenograft tissue more closely reflects the *in vivo* state compared to the HSE cultures with restored terminal differentiation, cell-cell adhesion, and partially restricted basal programs.

### Xenograft HSEs contain two distinct transcriptional trajectories

To characterize how the XENO clusters influence the keratinocyte’s differentiation trajectory, we employed pseudotime analysis overlayed onto the UMAP of the integrated *in vivo* and xenograft epithelial cells and found that xenografted keratinocytes likely follow two distinct transcriptional trajectories from basal to granular cells (**Figure 5A-B**). The XENO states are highly stable, along with the BAS-III state, whereas the other BAS, SPN, and GRN states are more unstable in the xenograft compared to their *in vivo* counterpart (**Figure 5C**). The inferred trajectory showed a progression from least differentiated to most differentiated for the xenograft-unique cell clusters, with progression from the highest *COL17A1+* state (XENO-4) to increasing *SBSN+* expression (XENO-1, XENO-2, and XENO-3) (**Figure 5D**). The splicing kinetics further supports two distinct differentiation trajectories, a BAS-SPN-GRN and a BAS-XENO-GRN trajectory possessing uniform velocity streams flowing from one state to the next (**Figure 5E**). The abundance of the XENO cluster cells (**Figure 4D**) suggests that the BAS-XENO-GRN differentiation trajectory is more favored in the xenograft.

**Figure 5.**
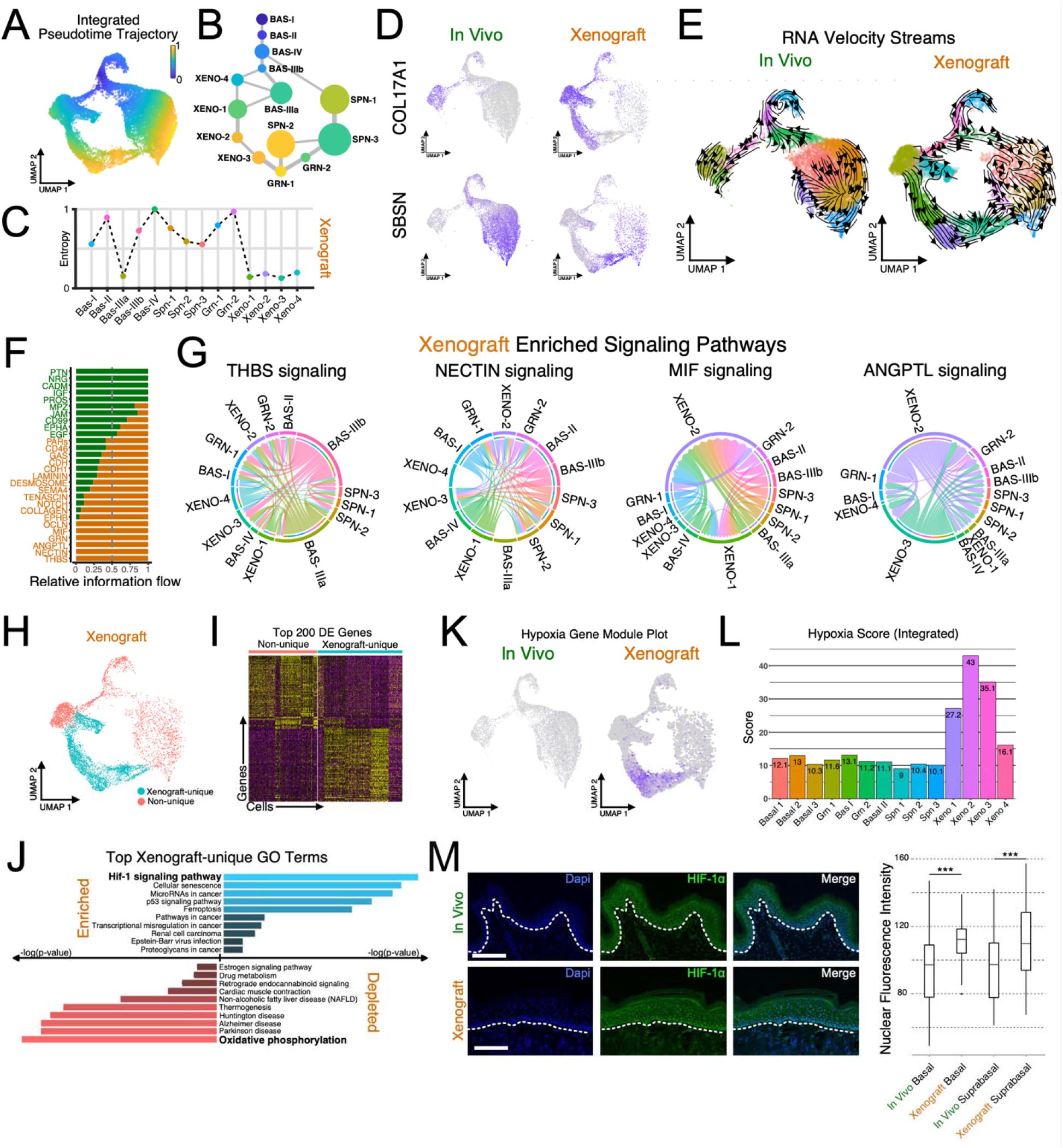
Hypoxia-driven transcriptional changes are observed in xenografts. **A)** Pseudotime inference and **B)** cell lineage diagram of epidermal keratinocytes from the integrated *in vivo* and xenograft datasets. Edge weights denote probability of transition to each cluster. Dot size denotes number of cells. **C)** Quantification of Cellular Entropy (*ξ*) using the **R** package SoptSC. **D)** Feature plots showing *SBSN* and *COL17A1,* marking differentiated and undifferentiated keratinocytes, respectively. **E)** Splicing kinetics depicted as RNA velocity streams calculated using the python package scVelo. **F-G)** Significant cell-cell communication networks inferred using the R package, CellChat. **H)** Metaclustering of xenograft cells into xenograft-unique and non-unique cohorts. **I)** Heatmap showing the top 200 differentially expressed (DE) genes between the two metaclusters. X-axis represent cells from the xenograft dataset and y-axis are DE genes. Yellow represents a relatively higher expression while purple represents relatively low expression. **J)** GO analysis of the top DE genes shown in panel I. Blue bars indicate biological processes upregulated in xenograft-unique cells; red bars indicate biological process downregulated in xenograft-unique cells. **K)** Feature plots showing expression of a hypoxia gene module consisting of 34 hypoxia-related genes. **L)** Hypoxia score, calculated as the sum of the average expression of each gene in the hypoxia gene module. Number in bar represents percentage. **M)** Immunostaining of HIF1-α in human neonatal epidermis and xenograft tissue. Quantification of nuclear HIF1-α stain is shown to the right. Significance was determined by unpaired two-tailed *t* test. *p < 0.001.

When we compare the relative information flow for the xenograft and *in vivo* datasets for each significant imputed pathway, several pathways show exclusive enrichment in the xenograft (OCLN, MIF, GRN, ANGPTL, NECTIN, and THBS) as well as the *in vivo* (PTN, NRG, CADM, IGF, PROS) datasets (**Figure 5F, Supplemental Tables 4-5**). All of the pathways that are unique to the xenograft are also present in at least one of the HSE cultures (**Supplemental Tables 2-3**). Although their functional roles within the HSE cultures are unclear, their known roles in skin biology suggest significant remodeling of the tissue and the extracellular environment. THBS signaling mainly originates in the BAS-III, XENO-1, and XENO-4 clusters, whereas ANGPTL signaling mainly originates in XENO-2 and XENO-3 clusters (**Figure 5G**), and both are known to promote angiogenesis (Lawler and Lawler, 2012; Oike et al., 2004), suggesting that the XENO clusters within the xenograft tissue may be hypoxic. NECTIN signaling shows promiscuous signaling throughout each cluster (**Figure 5G**), which is to be expected given its role in cell adhesion and skin morphogenesis (Okabe et al., 2004). The MIF signaling pathway largely signals to XENO-1 and XENO-2 clusters (**Figure 5G**) and has been shown to be upregulated during wound healing in mice (Gilliver et al., 2011).

### Hypoxia likely drives transcriptome-wide changes in xenograft-unique cells

As HSE-unique signaling pathways indicate significant tissue remodeling including enrichment for pathways that promote angiogenesis, we hypothesized that hypoxia may be driving the alternative differentiation trajectory in the XENO clusters. To explore this possibility, xenograft cells were metaclustered into two groups, xenograft-unique (XENO-1 through XENO-4) and non-unique clusters (BAS, SPN, and GRN) (**Figure 5H**). The xenograft-unique and non-unique metaclusters showed unique gene expression signatures (**Figure 5I**) and gene ontology analysis was performed on the top 100 marker genes for each metacluster using the KEGG 2019 Pathway database (**Figure 5J**). The most significantly enriched term for the xenograft-unique metacluster identified the HIF-1 signaling pathway, whereas the most significantly depleted pathway was oxidative phosphorylation (**Figure 5J**), which been shown to be down-regulated in response to hypoxia (Rodríguez-Enríquez et al., 2010). To explore this relationship further, we created a hypoxia gene module using Seurat’s gene module function that included a manually curated list of 34 genes that have been experimentally shown to be upregulated in response to hypoxia and/or possess a hypoxia response element in the promoter region (Kur-Piotrowska et al., 2018; Mieremet et al., 2019; Mole et al., 2009; Ngo et al., 2007) (**Supplemental Table 6**). The hypoxia gene module showed enhanced gene expression in the xenograft-unique metacluster with particular enrichment in the XENO-1, XENO-2, and XENO-3 clusters (**Figure 5K**), suggesting that the xenograft tissue is under hypoxic conditions. Using the sum of the hypoxia gene module expression to create a hypoxia score, XENO clusters 1-3 show high enrichment of hypoxia-related genes (**Figure 5L**). To validate the gene expression module and hypoxia score, we immunostained the xenografted HSE for the transcription factor HIF1A and found nuclear HIF1A expression is significantly higher in the xenografts than the *in vivo* tissues (**Figure 5M**), suggesting that hypoxia is causing widespread transcriptional changes to the xenografted keratinocytes.

## DISCUSSION

Human skin equivalents have long served as models of human IFE in place of murine skin (Augustin et al., 1997; Bell et al., 1983; El Ghalbzouri et al., 2009; Gu et al., 2020). We have shown that basal cell heterogeneity in our organoids fully mimics *in vivo* basal cell heterogeneity during homeostasis, with most of the differentiated states also present. However, HSE cultures exhibited signaling patterns characteristic of EMT events, contained organoid-unique cell states not found in *in vivo* neonatal epidermis where the cells were initially isolated, and showed differentiation abnormalities. Xenografting GelHSE cultures onto NSG mice rescued many of the defects in HSE cultures, but harbored xenograft-unique cell states likely driven by hypoxic conditions. These hypoxic conditions would likely last until the transplanted tissues reach homeostasis and wound repair pathways cease. For instance, wounding keratins KRT6/KRT16 were expressed in the grafted region at both days 16 and 37 in HSEs transplanted onto humans, with their expression disappearing a year after transplantation (Ojeh et al., 2017). Similarly, KRT14 was expressed in all layers of the epidermis until a year post-grafting where it resumed a normal basal layer expression, suggesting that the tissue did not reach homeostasis until a year post-grafting (Ojeh et al., 2017). However, transplantation of HSEs onto burn patients or recent transplantation of HSEs to cure junctional epidermolysis bullosa demonstrate their clinical importance and remains the gold standard (Hirsch et al., 2017).

Although basal cell heterogeneity was intact in the HSE and xenograft tissues, the proportion of BAS-III cells were enriched and BAS-IV cells were depleted compared to the *in vivo* state. BAS-III cells typically sit atop the rete ridges *in vivo,* whereas BAS-IV cells lie at the bottom of rete ridges (Wang et al., 2020b). However, this spatial environment is lost in the HSEs as the devitalized human dermis tends to flatten out during processing (**Figure 1B**), suggesting that spatial positioning may be important to specify the correct proportion of BAS-III-to-BAS-IV cells. The BAS-III state also shows more stability than BAS-IV, and BAS-III transcripts are retained throughout most of the other cellular states, suggesting the BAS-III program is not sufficiently shut down and may be the underlying cause of the differentiation defects seen in the SPN and GRN layers. Inappropriate signals from the dermis may also be the cause of the BAS defects. Although basal cells in both HSEs expressed canonical basal layer markers, they also expressed EMT-specific genes such as *VIM, LAMC2,* and *LGALS1.* The expression of these genes were higher in the GelHSE but were still present in FibHSE, suggesting that while Matrigel may be enhancing EMT-like programs, replacing Matrigel with primary human dermal cells is not sufficient to induce the appropriate *in vivo* expression programs and may be due to the culture media. Our results also suggest that HSEs may represent a wound regeneration or development model due to their EMT features and inappropriate expression of KRT14 (Haensel and Dai, 2018).

We identified two keratinocyte populations unique to the *in vitro* HSEs, one shared (HSE-2) between both HSEs and one specific (HSE-1) to the GelHSE. It is unclear where the shared population is located due to a limited number of marker genes, however, we suspect that HSE-2 is likely located closer to the cornified layer. As the HSE-2 cluster is placed after the HSE-1 population in pseudotime and that PSCA+ HSE-1 cells are located in the outermost HSE layers, the most likely localization of HSE-2 cells would be in a layer distal to the HSE-1 keratinocytes. Curiously, Psca expression occurs in the outermost layers of murine skin epithelium during E15-E17 (Ross et al., 2001). During this time, the outermost epithelial layer of the murine epidermis is periderm, which forms during stratification at E11.5 and disaggregates between E16-E17 when barrier formation occurs (Hammond et al., 2019). The periderm temporally expresses different marker genes as the epidermis differentiates, such as Krt17 during early stages and Krt6 during later stages (Hammond et al., 2019). *Psca* is upregulated in E18.5 murine epidermis of *Cyp26b1 −/−* mice which retains the periderm, suggesting *Psca* may be a marker gene of periderm at later stages. Taken together, this data suggests that primary keratinocytes from newborn epidermal tissue may retain enough plasticity to differentiate into prenatal cell types that are no longer found postnatally.

The presence of abnormal cell states and altered differentiation patterns in organoid cultures have been observed in a variety of tissues (Corrò et al., 2020), including skin (Atwood and Plikus, 2021), using more conventional methods. Matrigel is used in the majority of organoid systems (Corrò et al., 2020) and more than likely induces similar effects to those observed here. Recent papers using scRNA-seq to characterize organoid cultures of other tissue types have also identified abnormal cell populations present in their organoid cultures. For instance, melanoma-like, neuronal-like, and muscle-like cells were found using scRNA-seq of kidney organoids (Subramanian et al., 2019), which were consistent with previous observations using conventional methods in this system. scRNA-seq analysis of human intestinal and brain organoids used random forest classifiers to identify the cell types in their organoid cultures (Fujii et al., 2018; Velasco et al., 2019), however, doing so excludes the possibility of classifying cells as anything other than predefined types. This is true of any supervised machine learning algorithm and can be misleading when examining cellular heterogeneity.

Despite the transcriptional and molecular differences we see in HSE organoid cultures, they still are attractive systems for investigative dermatology and are superior to two-dimensional tissue culture of primary keratinocytes. Both HSE culture conditions form fully stratified tissues, generate most of the *in vivo* cellular states, and largely reach homeostatic conditions after transplantation. Although xenografted HSEs are still utilizing wound repair programs 24 days post engraftment, allowing more time for the graft to heal would presumably return it to a fully homeostatic state. Potential ways to improve HSEs to more faithfully mimic *in vivo* skin could include the addition of cell types such as Langerhans cells, melanocytes, endothelial cells, and other immune cells. Altering culturing conditions or bioengineering 3D scaffolds may also help restrict basal and terminal differentiating programs to their proper cellular states.

## METHODS

### Ethics statements

Human clinical studies were approved by the Ethics Committee of the University of California, Irvine. All human studies were performed in strict adherence to the Institutional Review Board (IRB) guidelines of the University of California, Irvine (2009-7083). We have obtained informed consent from all participants.

### Preparation of devitalized dermis

Cadaver human skin was acquired from the New York Firefighters Skin Bank (New York, New York, USA). Upon arrival at UC Irvine, the skin was allowed to thaw in a biosafety cabinet. Skin was then placed into PBS supplemented with 4X Pen/Strep, shaken vigorously for 5 minutes, and transferred to PBS supplemented with 4X Pen/Strep. This step was repeated two additional times. The skin was then placed into a 37°C incubator for 2 weeks. The epidermis was removed from the dermis using sterile watchmaker forceps. The dermis was washed 3 times in PBS supplemented with 4X Pen/Strep with vigorous shaking. The dermis was then stored in PBS supplemented with 4X Pen/Strep at 4°C until needed.

### Primary cell isolation

Discarded and de-identified neonatal foreskins were collected during routine circumcision from UC Irvine Medical Center (Orange, CA, US). The samples were either processed for histological staining, single cell RNA-sequencing, or primary culture. No personal information was collected for this study. For primary cell isolation, fat from discarded and deidentified neonatal foreskins were removed using forceps and scissors and incubated with dispase epidermis side up for 2 hours at 37°C. The epidermis was peeled from the dermis, cut into fine pieces, and incubated in 0.25% Trypsin-EDTA for 15 minutes at 37°C and quenched with chelated FBS. Cells were passed through a 40μm filter, centrifuged at 1500rpm for 5 minutes, and the pellet resuspended in Keratinocyte Serum Free Media supplemented with Epidermal Growth Factor 1-53 and Bovine Pituitary Extract (Life Technologies; 17005042). Cells were either live/dead sorted using SYTOX Blue Dead Cell Stain (ThermoFisher; S34857) for single cell RNA-sequencing or incubated at 37°C for culture.

### Cell sorting

Following isolation, cells were resuspended in PBS free of Ca^2+^ and Mg^2+^ and 1% BSA and stained with SYTOX Blue Dead Cell Stain (ThermoFisher; S34857). Samples were bulk sorted at 4°C on a BD FACSAria Fusion using a 100μm nozzle (20 PSI) at a flow rate of 2.0 with a maximum threshold of 3000 events/sec. Following exclusion of debris and singlet/doublet discrimination, cells were gated on viability for downstream scRNA-seq.

### Human skin equivalent organoid culture

Primary human keratinocytes were cultured in Keratinocyte Serum Free Media supplemented with Epidermal Growth Factor 1-53 and Bovine Pituitary Extract (Life Technologies; 17005042). Generation of organotypic skin cultures were performed as described in Li and Sen, 2015. Briefly, ~500K control cells were seeded on devitalized human dermis and raised to an air/liquid interface to induce differentiation and stratification over the indicated number of days with culture changes every two days. Prior to seeding keratinocytes, either Matrigel was applied to the underside of the devitalized dermis or primary human dermal fibroblasts were centrifuged into the devitalized dermis. To measure changes from EGF supplementation, culture media was switch to Keratinocyte Serum Free Media supplemented with Bovine Pituitary Extract and variable concentrations of Epidermal Growth Factor 1-53 (Life Technologies; 17005042) after one week for one additional week of culturing.

### Human skin equivalent xenograft model

Human neonatal epidermal keratinocytes (Thermo Fisher Scientific; C0015C) were maintained in Epilife media (Thermo Fisher: MEPI500CA) supplemented with HGKS (Thermo Fisher: S0015). To generate skin equivalents, 10^6 cells were seeded onto devitalized human dermis and maintained in an air-liquid interface for 7 days. Stratified epithelial tissue was then grafted onto 12-14 week old female NOD scid gamma mice (Jackson Laboratory: 005557). Bandages and sutures were removed 2 weeks after surgery and healthy grafts were harvested 10 days later.

### Droplet-enabled single cell RNA-sequencing and processing

Cell counting, suspension, GEM generation, barcoding, post GEM-RT cleanup, cDNA amplification, library preparation, quality control, and sequencing was performed at the Genomics High Throughput Sequencing Facility at the University of California, Irvine. Transcripts were mapped to the human reference genome (GRCh38) using Cell Ranger Version 2.1.0.

### Histology and immunohistochemistry

Frozen tissue sections (10μm) were fixed with 4% PFA in PBS for 15 minutes. Following fixation, tissue sections were stained with Hematoxylin and Eosin following standard procedures. Sections were stained with Gill’s III (Fisher Scientific; 22050203) for 5 minutes and Eosin-Y (Fisher Scientific; 22050197) for 1 minute. Tissue sections were visualized under a light microscope under 10x objective lens after mounting with Permount mounting media (Fisher Scientific; SP15-100). For immunostaining, tissue sections were fixed with 4% PFA in PBS for 15 minutes. 10% BSA in PBS was used for blocking. Following blocking, 5% BSA and 0.1% Triton X-100 in PBS was used for permeabilization. The following antibodies were used: chicken anti-KRT14 (1:500; BioLegend; SIG-3476), rabbit anti-KI67 (1:500; Abcam; ab15580), rabbit anti-COL17A1 (1:100; One World Labs; ap9099c), rabbit anti-KRT19 (1:250; Cell signaling; 13092), mouse anti-KRT15 (1:500; Santa Cruz; sc-47697), rabbit anti-VIM (1:500; Cell Signaling; D21H3), mouse anti-PSCA (1:500; Santa Cruz; sc-80654), mouse anti-FLG (1:500; Santa Cruz; sc-66192), mouse anti-DSG1 (1:500; Santa Cruz; sc-137164), and rabbit anti-LOR (1:500; Abcam; ab85679). Secondary antibodies included Alexa Fluor 488 (1:500; Jackson ImmunoResearch; 715-545-150, 711-545-152) and Cy3 AffiniPure (1:500; Jackson ImmunoResearch; 711-165-152, 111-165-003). Slides were mounted with Prolong Diamond Antifade Mountant containing DAPI (Molecular Probes; P36962). Confocal images were acquired at room temperature on a Zeiss LSM700 laser scanning microscope with Plan-Apochromat 20x objective or 40x and 63x oil immersion objectives. Images were arranged with ImageJ, Affinity Photo, and Affinity Designer.

### Quality control metrics post-Cell Ranger assessment

For downstream analyses, we kept cells which met the following filtering criteria per biological replicate per condition: >200 and <10% mitochondrial gene expression. Genes that were expressed in less than 3 cells were excluded. Data was normalized with a scale factor of 10,000.

### Data availability

The authors declare that all data supporting the findings of this study are available within the article and its supplementary information files or from the corresponding author upon reasonable request. The datasets generated during the current study have been deposited at the GEO database under accession code GSE190695 [https://www.ncbi.nlm.nih.gov/geo/query/acc.cgi?acc=GSE190695].

## ACKNOWLEDGEMENTS

S.X.A. is supported by NIH grant R01CA237563 and NSF grant 2134916. Q.N. is supported by NIH grants U01AR07315 and R01GM123731, an NSF grant DMS11763272, and a Simons Foundation grant 594598. The authors wish to acknowledge the support of the Chao Family Comprehensive Cancer Center Genomics High-Throughput Facility and Optical Biology Core Shared Resource, supported by the National Cancer Institute of the NIH under award number P30CA062203, and the UCI Skin Biology Resource Center supported by the National Institute of Arthritis and Musculoskeletal and Skin Diseases under award number P30AR075047. We also thank Jennifer M. Bates and the UCI Institute for Immunology Flow Cytometry Core Facility for help with cell sorting and Bryan K. Sun for comments on the manuscript.

## AUTHOR CONTRIBUTIONS

S.X.A. and A.R.S. conceived the project; S.X.A. supervised research; A.R.S. generated and analyzed scRNA-seq libraries; S.W. performed SoptSC for lineage and entropy analysis; A.R.S., S.D.N., and G.E.L. performed imaging experiments; J.L. and G.L.S. performed xenograft experiments; A.R.S., S.W., Q.N., and S.X.A. analyzed and interpreted data; A.R.S. and S.X.A. wrote the manuscript. All authors analyzed and discussed the results and commented on the manuscript.

---

**Supplemental Figure 1.**
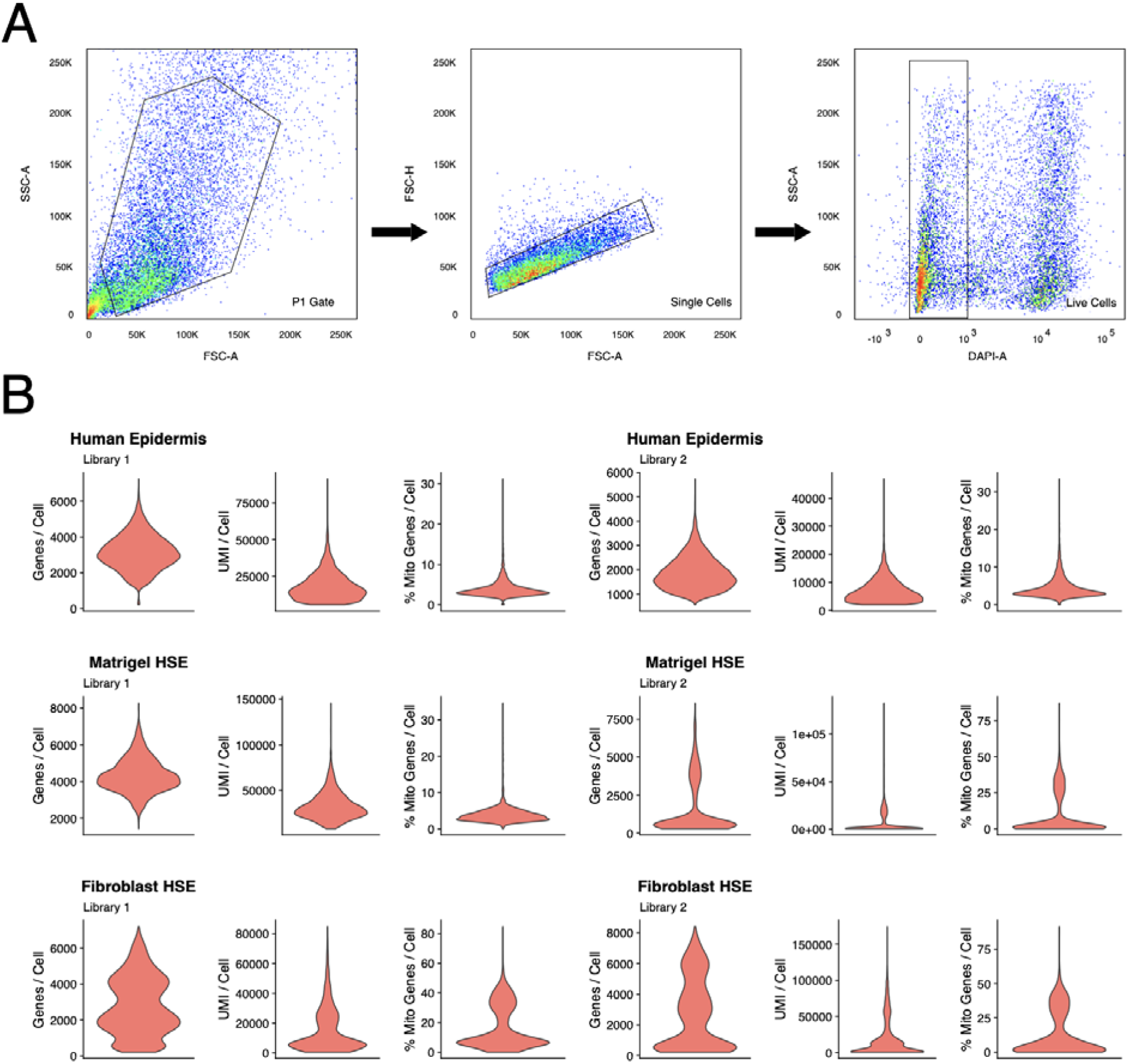
FACS strategy and quality control metrics. **A)** Schematic of FACS strategy for physical sorting of live cells from human skin equivalents. SSC – A denotes side scatter area. FSC – A denotes forward scatter area. FSC – H denotes forward scatter height. FITC – A denotes dead cells using SYTOX Blue. **B)** Violin plots showing genes per cell, percent mitochondrial (mito) genes per cell, and unique molecular identifiers (UMI) per cell for each *in vivo* (human epidermis), Matrigel HSE, and Fibroblast HSE libraries prior to quality control filtering.

**Supplemental Figure 2.**
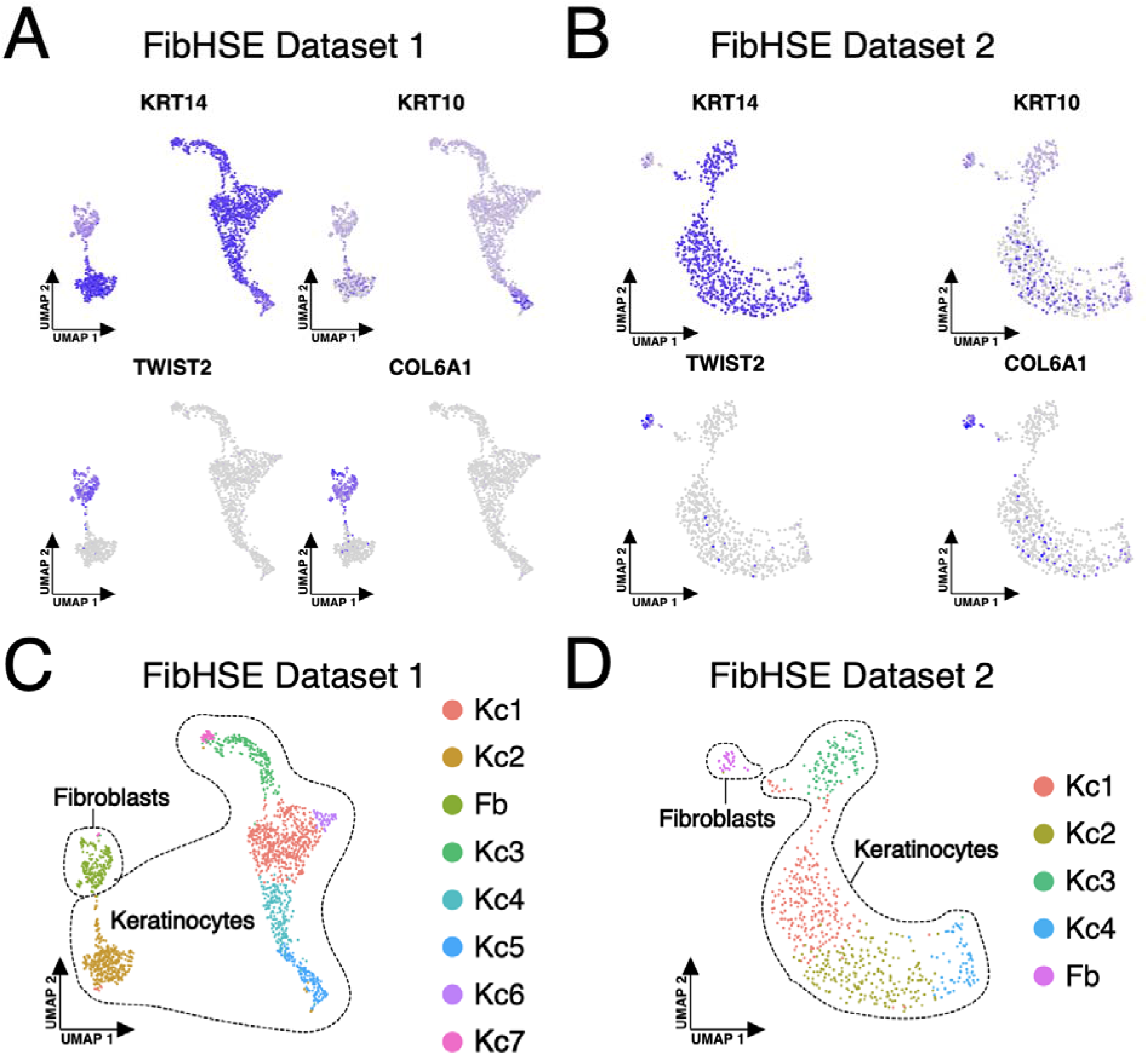
Identification of fibroblast populations in FibHSE datasets. Feature plots showing the expression of keratinocyte markers *KRT14* and *KRT10* and fibrobla**st** markers *TWIST2* and *COL6A1* for **A)** FibHSE dataset 1 and **B)** FibHSE dataset 2. Clustering **of** single cells isolated from **C)** FibHSE dataset 1 and **D)** FibHSE dataset 2 libraries display**ed** using UMAP embedding. Keratinocyte and fibroblast cell populations are outlined.

**Supplemental Figure 3.**
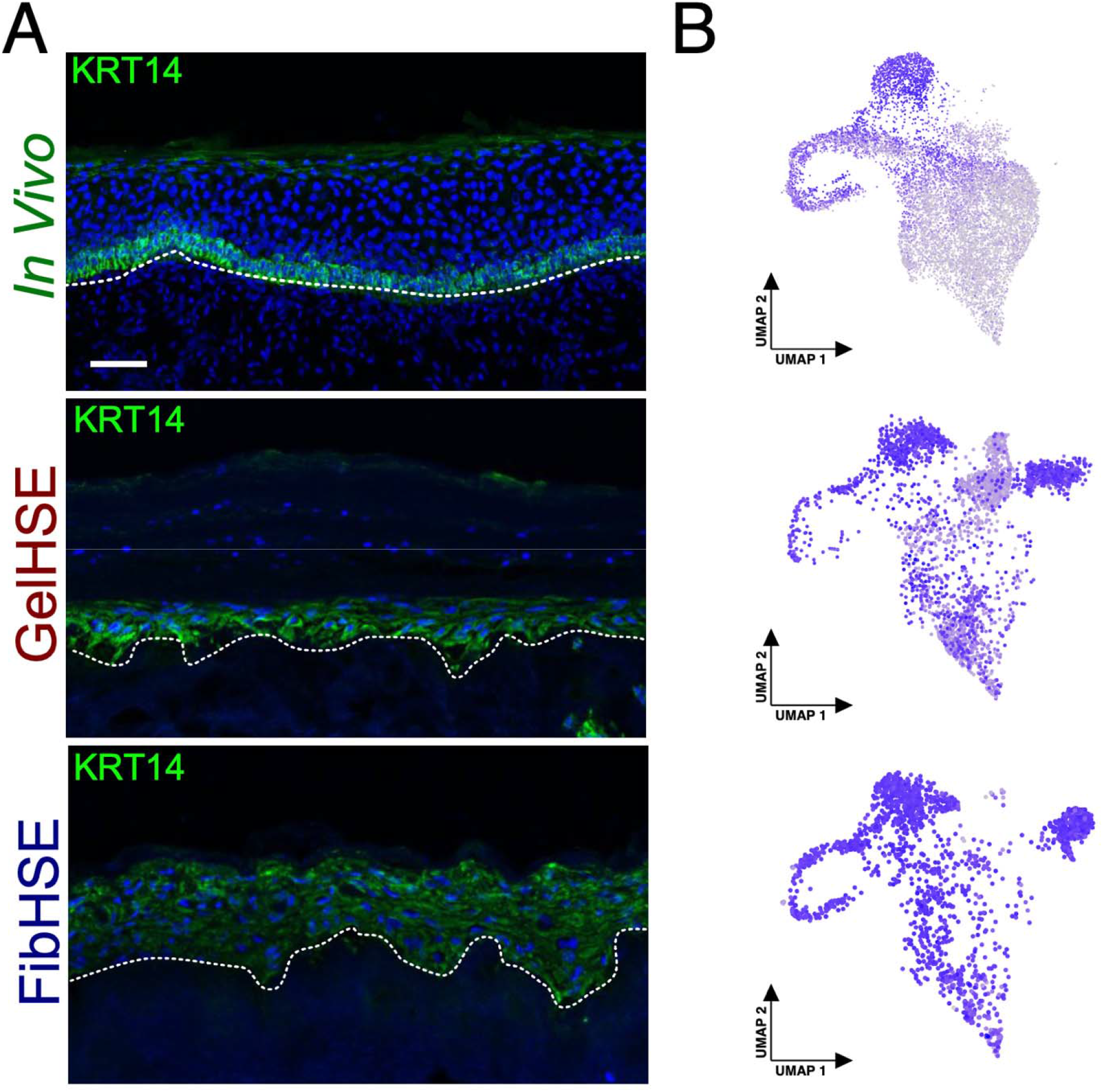
KRT14 expression in neonatal and HSE organoids. **A)** Immunostaining of KRT14 in human neonatal skin (top), day 28 GelHSE (middle), and day **28** FibHSE (bottom). **B)** Feature plots showing the RNA expression of *KRT14* for each sample type shown in panel A.

**Supplemental Figure 4.**
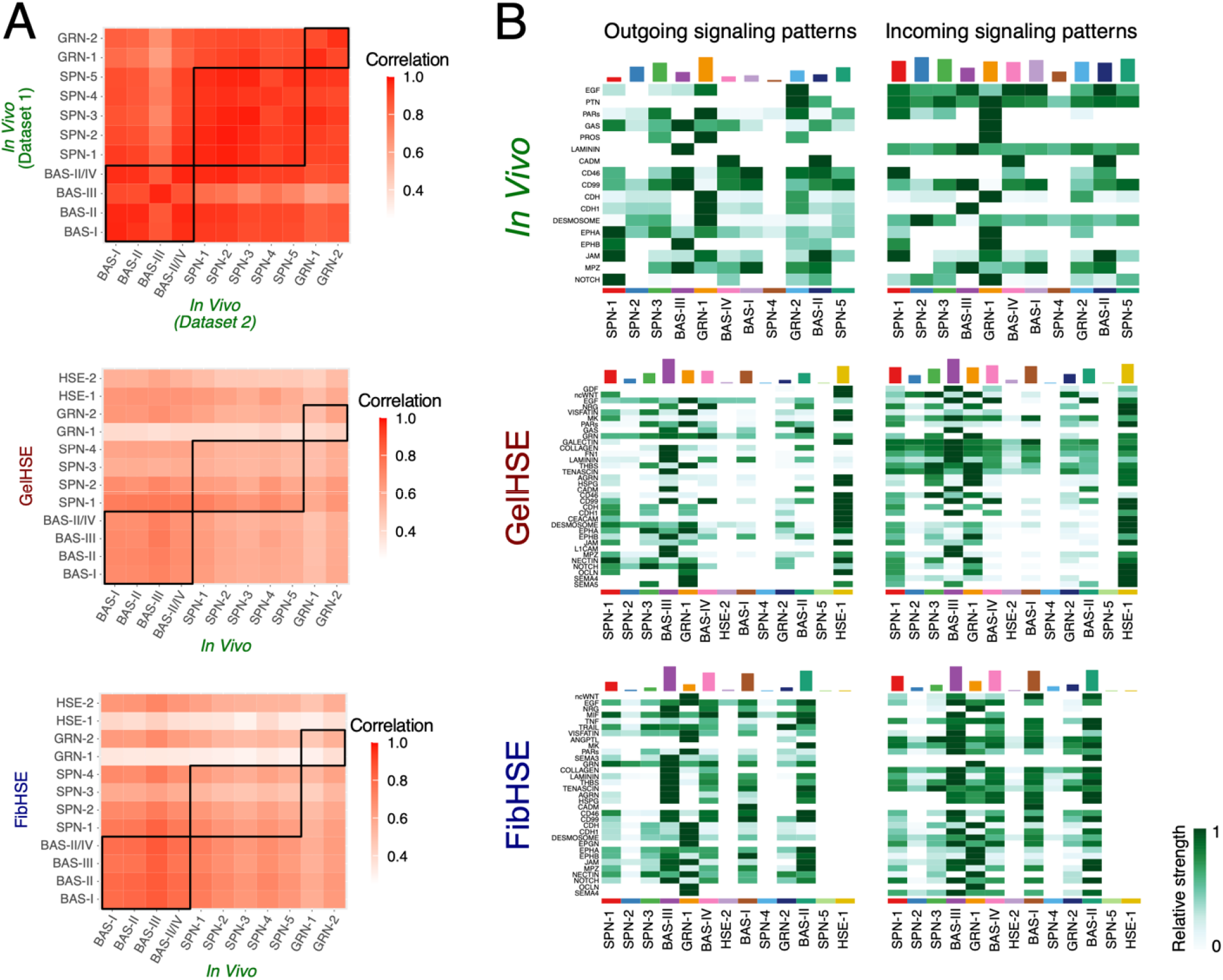
Pearson correlation and signaling analysis. **A)** Pearson correlation between average RNA expression of each cluster from the indicated datasets. **B)** Heatmap showing the relative strength of outgoing and incoming signals for all significant signaling pathways for the *in vivo,* GelHSE, and FibHSE datasets.

**Supplemental Figure 5.**
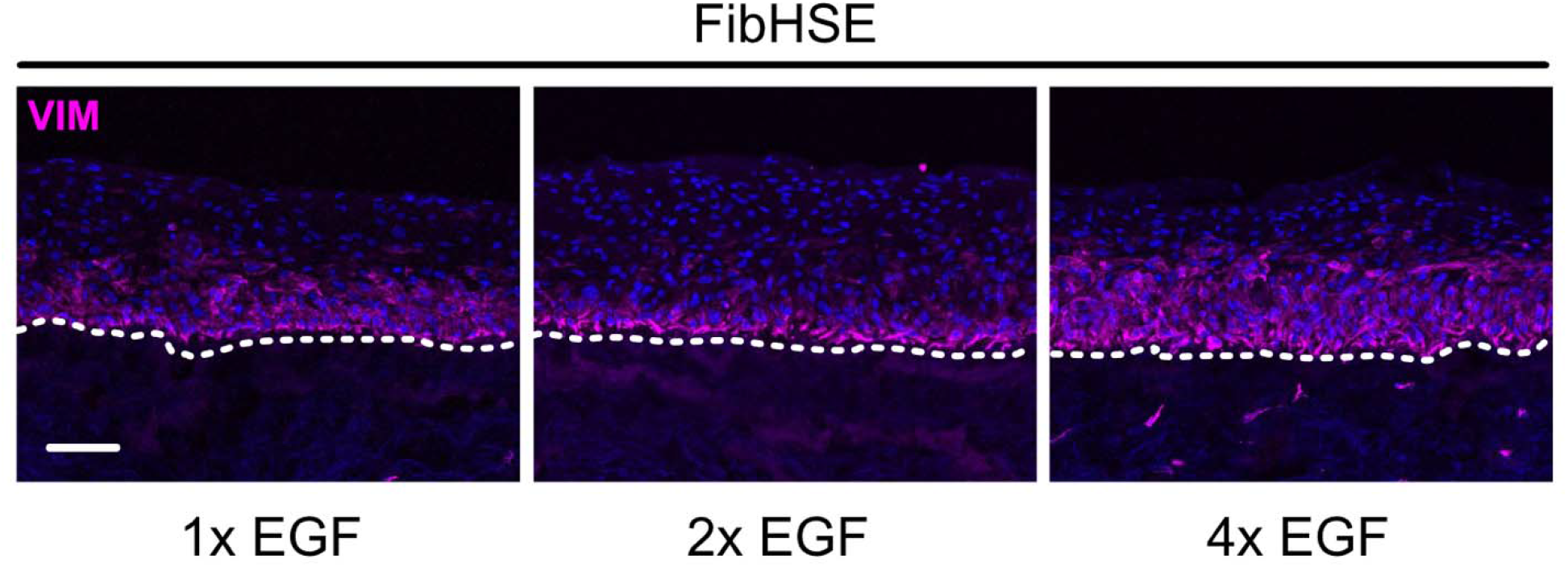
VIM immunostaining of day 14 FibHSEs supplemented with 1x (Left), 2x (middle), and 4x (right) EGF. Scale bar = 100 μm.

**Supplemental Figure 6.**
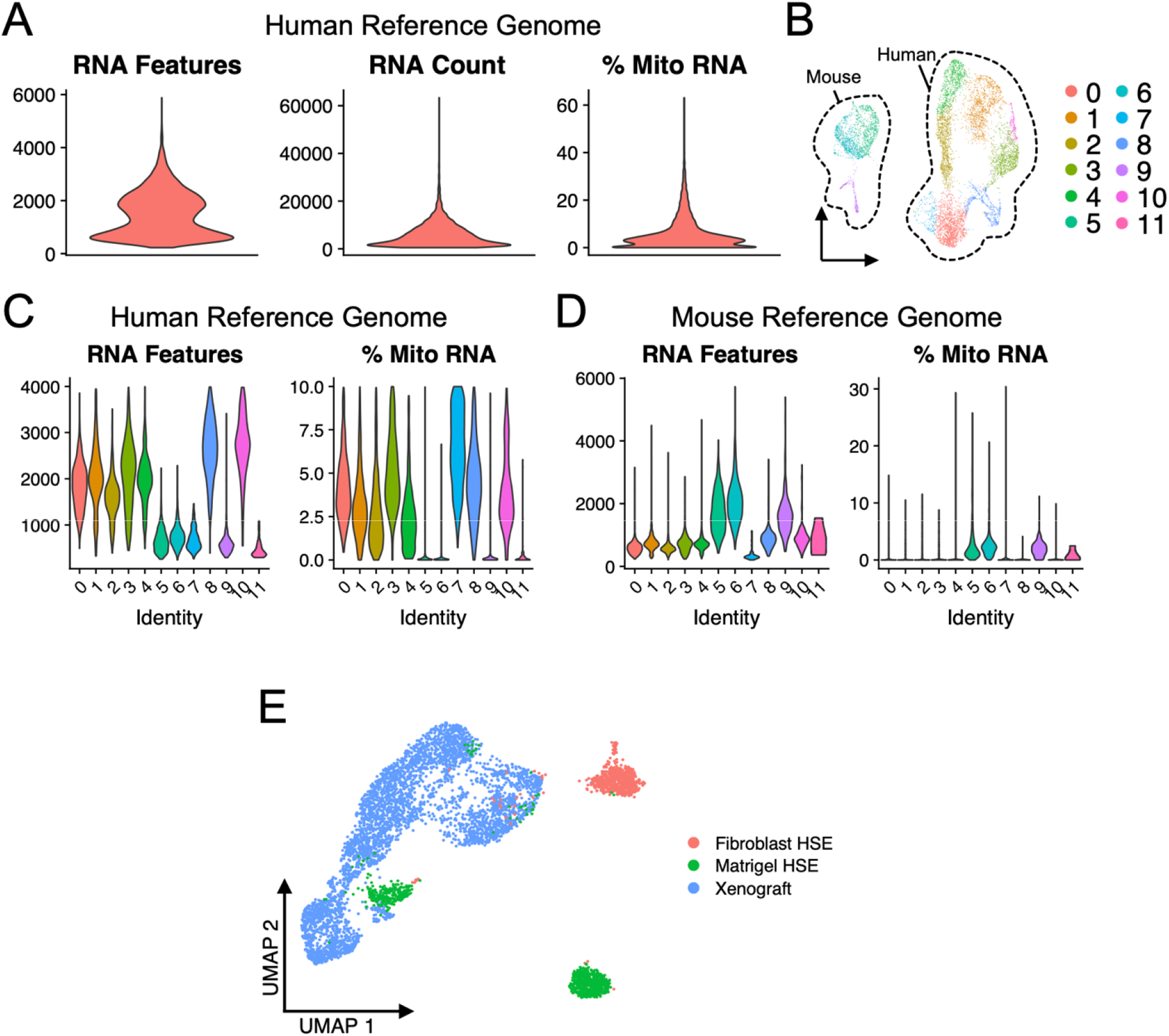
Quality control metrics of xenografted HSE dataset. **A)** Violin plots showing genes per cell, unique molecular identifiers (UMI) per cell, and percent mitochondrial (mito) genes per cell for the xenograft dataset aligned to the GRCh38 human reference genome. **B)** Clustering of single cells isolated from xenograft library displayed using UMAP embedding. Mouse cells and human cells are outline by dashed lines and labeled. **C-D)** Violin plots showing genes per cell and percent mito genes per cell for the xenograft dataset after clustering and aligning to the GRCh38 human reference genome **(C)** and mm10 mouse reference genome **(D)**. **E)** Integration and clustering of HSE-unique populations, HSE-1 and HSE-2, along with xenograft-unique populations, XENO-1 through −4, with sample type labels superimposed onto UMAP embedding.

**Supplemental Figure 7.**
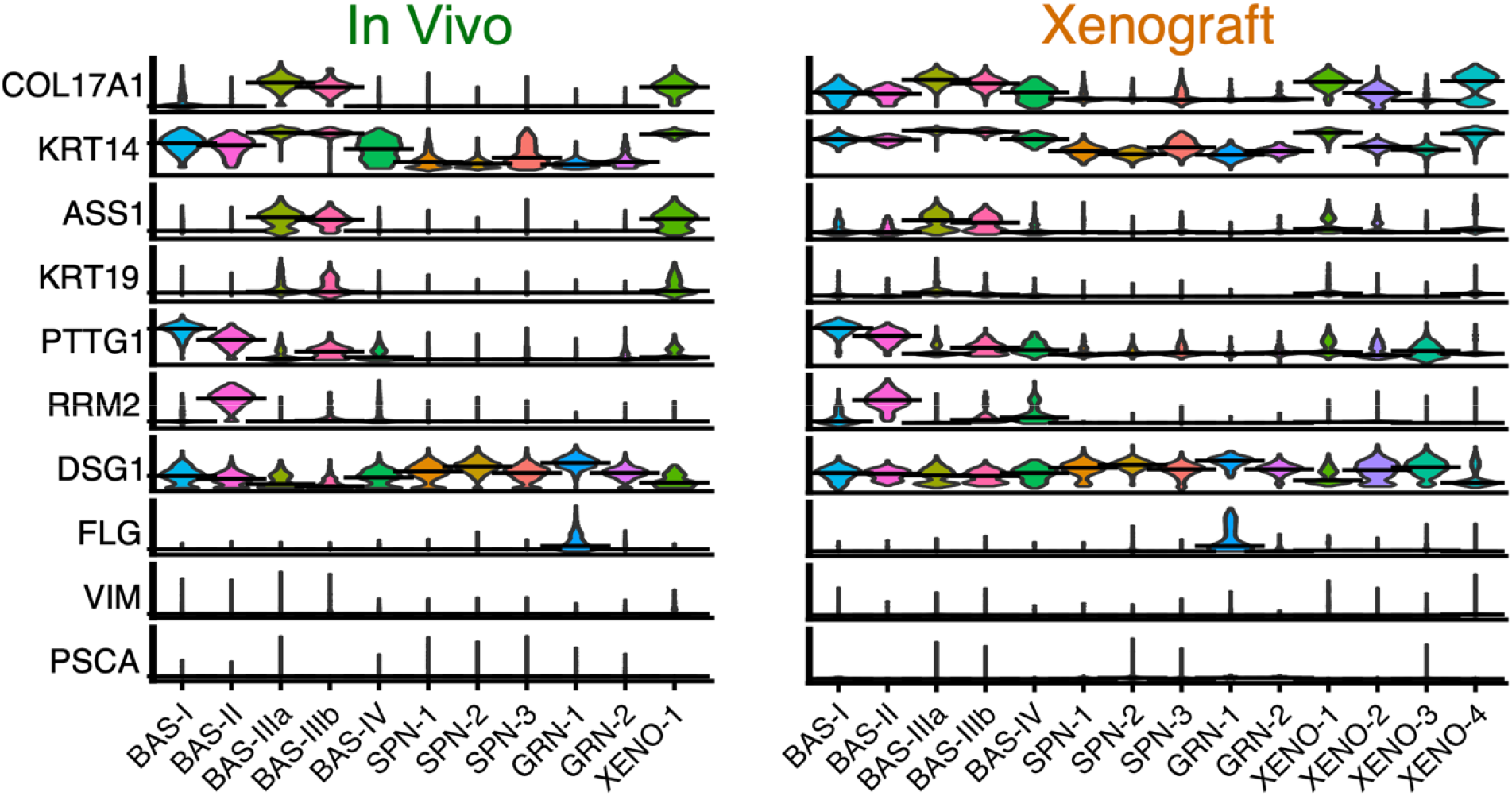
Differentially expressed genes in xenograft dataset. Violin plots showing RNA expression of the indicated genes in each cluster of the in vivo or xenograft datasets.

